# Development and validation of an ultra-low-cost, open source normothermic *ex vivo* organ perfusion platform

**DOI:** 10.1101/2025.10.27.684886

**Authors:** Heiko Yang, Nicholas Higgins, Simon Chu, Jean Lee, Natalie Meyer, Keith Hansen, Mustafa Saeed, Ryan Ferreira, Thomas A. Sorrentino, Jorge Mena, Pablo Suarez, Feres Camargo Maluf, Wilson Sui, Maria C. Velasquez, Uday Mann, Hillary Braun, Juan Du, Jacob Elmer, Tom Chi, Shuvo Roy, Alan Flake, James M. Gardner, Marshall Stoller

**Author notes:** These authors contributed equally (Heiko Yang, Nicholas Higgins, Simon Chu). Corresponding authors: Heiko Yang, MD, PhD 12631 E 17^th^ Ave, Mail Stop C312 Aurora, CO 80045 James M. Gardner, MD, PhD Health Sciences West, Box 0534 513 Parnassus Ave San Francisco, CA 94143.

## Abstract

**Background:** Normothermic *ex vivo* organ perfusion (NEVOP) promises to catalyze organ preservation, therapeutic discovery, and organ-specific disease modeling. Existing technology platforms remain inaccessible for research due to restricted access to commercial organ perfusion devices, high costs of both devices and proprietary consumables, and steep technical learning curves. Additionally, the available technology is not optimized to perfuse smaller organs such as the kidney.

**Methods:** To overcome these barriers, a custom NEVOP circuit was developed using recycled, repurposed, and low-cost components. Porcine kidneys and autologous blood were used to iteratively optimize circuit design. A porcine kidney autotransplantation protocol was adapted to evaluate *in vivo* kidney function after *ex vivo* perfusion. To pilot the flexibility of this system as a multi-organ platform for *ex vivo* human biology, non-transplantable human donor kidney, spleen, and pancreas specimens were stably perfused using human blood products and analyzed.

**Results:** An ultra low-cost NEVOP system engineered to perfuse porcine kidneys and diverse human organs (kidney, pancreas, and spleen) sustained viable organs for up to 24 hours with evidence of both function and viability. Key innovations included a parallel flow resistor to facilitate low-flow perfusion in non-heparinized organs and a containment bag with adjustable magnets to provide vascular stability and recycling of venous overflow. The circuit costs less than 1,500USD to construct, and porcine kidneys perfused for 24 hours on this platform demonstrated healthy *in vivo* function upon autotransplantation.

**Conclusions:** Custom NEVOP platforms constitute novel and potentially transformative research platforms which use low-cost and readily available materials. Paired with access to non-transplantable research organs from altruistic donors, this model provides a road map for investigators to advance biomedical discovery and human *ex vivo* biology.

## INTRODUCTION

Normothermic *ex vivo* organ perfusion (NEVOP), also known as normothermic machine perfusion, is a burgeoning field of biomedical exploration.^1–5^ The ability to sustain solid organs in a physiologic state outside the body using the principles of extracorporeal membranous oxygenation (ECMO) and cardiopulmonary bypass (CPB) promises to unlock numerous possibilities for organ preservation, organ-directed therapy, and disease modeling.^6–8^ This technology has already been used clinically in organ transplantation to prolong the viability of hearts, livers, and lungs, helping more donor organs reach suitable recipients.^9–13^ Recent studies have further demonstrated the therapeutic potential of this technology as an opportunity to therapeutically modify the biology of an organ.^14–17^

Despite the immense enthusiasm around NEVOP, few laboratories worldwide conduct this research. There are numerous barriers to entry. The cost of NEVOP devices currently ranges between tens to hundreds of thousands of US dollars per unit.^18^ The proprietary nature of these devices also limits their availability for academic research. Acquiring suitable reagents can be challenging for inexperienced investigators. Not only do organ perfusion experiments often rely on specialized reagents such as University of Wisconsin Solution, HTK Solution, or STEEN Solution, but they also require sourcing blood and organs from large laboratory animals or human donors. The steep technical learning curve also compounds the costs and logistical challenges associated with each experiment. Given these barriers, the activation energy required to enter the field of organ perfusion research and to maintain this line of inquiry can appear to be insurmountable.

To solve these problems, we developed a practical, ultra-low-cost approach to building a NEVOP system and attaining proficiency in organ perfusion. Initially, a normothermic *ex vivo* kidney perfusion (NEVKP) system was developed to address the need for a device tailored specifically to perfuse kidneys.^7^ As such, a circuit that included a mechanical pressure regulation system and a magnetized containment bag was engineered. Inexpensive, recycled, and repurposed materials were utilized to reduce costs during the initial learning curve. A strategy to perfuse nonheparinized porcine organs, which helped to reduce the cost of obtaining porcine blood and organs for perfusion research was undertaken. A porcine renal autotransplantation protocol to test *in vivo* kidney viability following prolonged NEVKP was adapted. Our perfusion system subsequently was adjusted to other human organs such as the pancreas and spleen, and functional and viability assessments were tailored to each organ.

## METHODS

### Circuit design

Three circuit design iterations were used in this study (Fig. 1A). Each circuit consisted of a peristaltic roller pump (Maquet or Cole-Parmer), a Capiox FX05 neonatal oxygenator (Terumo), and 0.25-inch Tygon tubing (Masterflex L/S 17) with polycarbonate straight and T-connectors (McMaster-Carr). Microclave needleless valves (ICU Medical Inc) were used for sample ports. Carbogen (95% O_2_, 5% CO_2_, Airgas) was delivered to the oxygenator using a rotameter (Snowate LZB-4W(B)). A digital blood pressure analyzer was used to monitor perfusion pressure (Digi-Med). The cost of all components is listed in Table 1.

**Figure 1.**
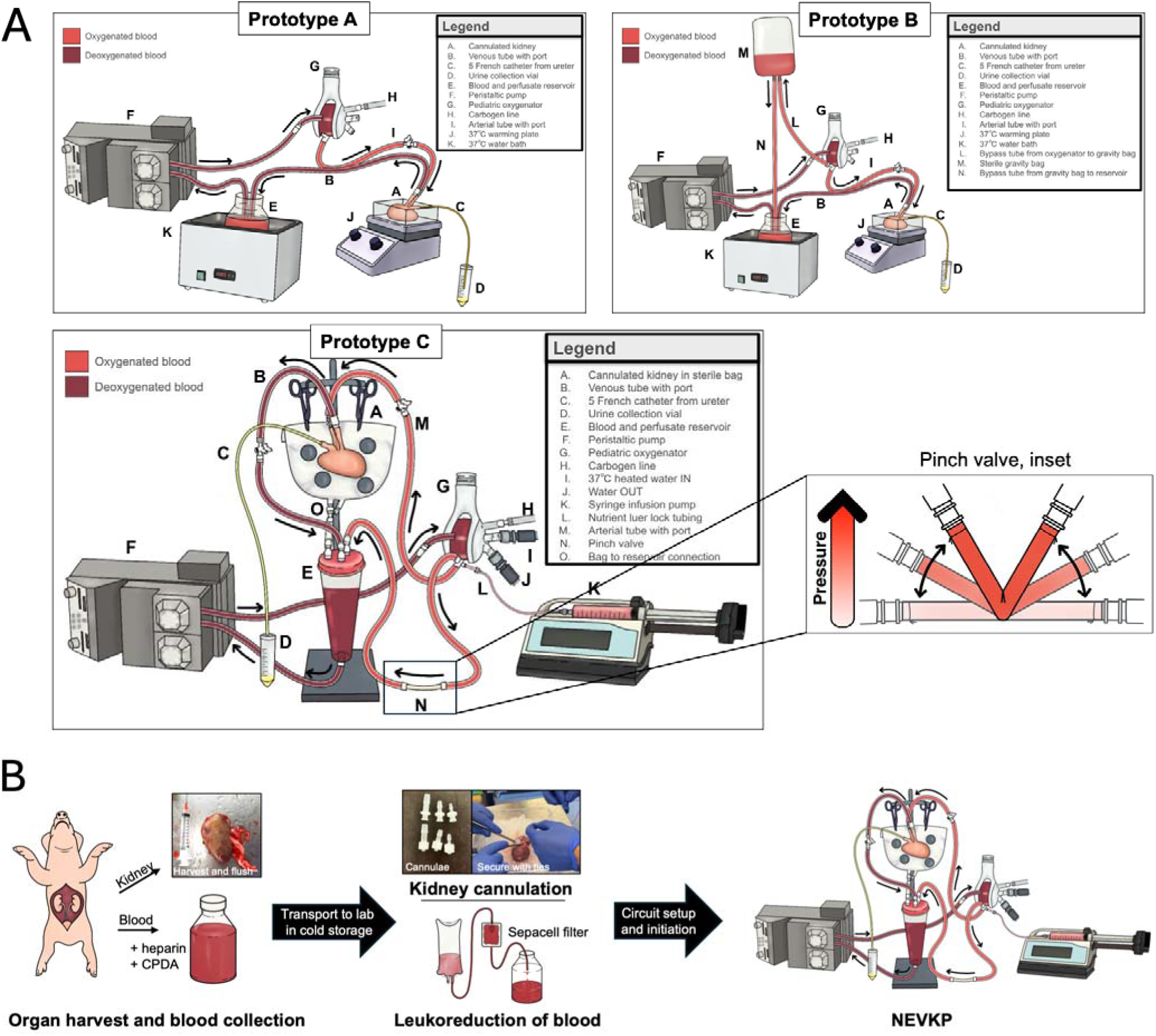
Blueprint for low-cost NEVKP. A. Three NEVKP circuits reflecting the chronologic evolution of our circuit design. B. Experimental workflow from harvesting tissue to initiating NEVKP.

**Table 1.** Core circuit components and availability for purchase. Prices updated as of August 2025.

| Component | Source | Per-unit cost | Quantity needed per circuit | Item cost per circuit | Reusable? |
| --- | --- | --- | --- | --- | --- |
| Masterflex® Precision Pump Tubing LS-17, 25 ft | Masterflex | \$187.20 | 10 ft | \$93.60 | Yes |
| LS 17 1/4 Luer Locks Male | Amazon | \$0.27 | 15 | \$4.00 | Yes |
| LS 17 1/4 Luer Locks Female | Amazon | \$0.33 | 15 | \$5.00 | Yes |
| 3-way Luer Lock Connectors | Amazon | \$1.59 | 2 | \$3.18 | Yes |
| MicroClave Neutral Connectors | Amazon | \$1.73 | 2 | \$3.46 | No |
| 1/4in Straight Connectors | Amazon | \$1.16 | 2 | \$2.32 | Yes |
| 1/4 to 3/8in Straight Connectors | Amazon | \$1.40 | 2 | \$2.80 | Yes |
| Peristaltic Pump | Kamoer / Amazon | \$200 | 2 | \$400 | Yes |
| Silicone Penrose Drains | Medline | \$4.04 | 1 | \$2.02 | Yes |
| Reservoir (Modified T225 Flask or neonatal suction canister ) | Thermo Fisher Scientific | \$10.00 | 1 | \$10.00 | Yes |
| Capiox FX05 Baby-FX Oxygenator | Terumo | \$725 | 1 | \$725 | Yes |
| 15mm Magnets | Amazon | \$0.62 | 12 | \$7.44 | Yes |
| 3 L Saline Bags | Baxter | \$27.00 | 1 | \$27.00 | No |
| Blood Pressure Analyzer | Digi-Med / Ebay | \$49.99 | 1 | \$49.99 | Yes |
| Gas rotameter | Ebay | \$21.30 | 1 | \$21.30 | Yes |
| | | | **Total cost** | \$1,357.11 | |

Prototype A: This initial circuit was designed with a pump, oxygenator, and kidney in series. A plastic 1 L bottle was used as the reservoir; warming was provided by placing the reservoir in a water bath and placing the kidney container over a hot plate set at 40 °C. The kidney was housed in an open polytetrafluoroethylene (PTFE) container filled with Plasma-Lyte (Baxter).

Prototype B: This subsequent design was like prototype A except for the addition of a bypass segment between the arterial recirculation port on the oxygenator to the reservoir. An empty, open-to-atmosphere 1 L bag was suspended ∼80-120cm above the kidney to assist in controlling perfusion pressure.

Prototype C: In this design, the gravity bag mechanism was replaced with a short segment of silicone Penrose tubing (0.25-inch diameter, Medline) that could be variably occluded to provide pressure control. A custom reservoir was created from a neonatal suction canister that was modified to include a drainage port at the bottom, four ports in the lid (venous, bypass, kidney bag, urine). Warming was accomplished by running warmed water through the oxygenator using either a standard peristaltic pump (Cole-Parmer) or an ECMO warmer (Cincinnati Sub-Zero). The cannulated kidney was housed in a plastic 3 L bag (Baxter) modified to include a drainage port using either the infusion or spike port. The kidney was secured in place within the bag with a series of 9mm x 3mm neodymium disc magnets (MIN CI). A small opening in the side was created to allow the ureteral catheter to exit. After securing the cannulas and ureter within the bag, the apparatus was suspended above the reservoir.

Of note, for prolonged NEVKP experiments (≥12 hours), the peristaltic pump was replaced with a centrifugal pump (Terumo) to reduce hemolysis.^19^

### Reuse of circuit components

During the circuit development phase, disposable items (oxygenators, tubing, connectors, reservoirs) were cleaned between experiments and reused to reduce cost. Each component was rinsed with deionized water, 10% bleach, Tergazyme (Alconox), and water again until clear. Prior to storage, oxygenators were dried overnight using compressed air.

### Nonheparinized porcine kidneys and autologous porcine blood

Nonheparinized porcine kidneys and autologous blood were primarily obtained from two sources: during necropsy for other investigators’ porcine experiments within our animal facilities or from a local slaughterhouse. In each case, organs and blood were obtained without charge.

In the case of necropsy animals, 1 L autologous whole blood was obtained upon exsanguination for euthanasia. CPDA-1 (150 mL per L) and heparin (5000 u per L) were added immediately to prevent clot formation. After cardiac death, kidneys were extracted from bilateral flank incisions and flushed immediately with at least 100 mL cold heparinized saline or Viaspan solution (Belzer) until effluent ran clear.

In the case of slaughterhouse animals, animals were electrically stunned and then immediately exsanguinated from the jugular vein. Similar to above, CPDA-1 (150 mL per L) and heparin (5000 u per L) were added immediately to the collected blood to prevent clot formation. Animals were then eviscerated within 10 minutes of exsanguination. The kidneys were identified within the entrails and dissected free. Once dissected, kidneys were flushed immediately with at least 100 mL cold heparinized saline or Viaspan solution (Belzer) until effluent ran clear.

After flushing, all organs were immersed in cold Viaspan solution for transportation back to the laboratory prior to NEVOP. Blood was subsequently leukoreduced using a Sepacell RS-2000 filter (Fenwal; 500 mL per filter) prior to use.

### Human donor organs and allogenic blood products

Human donor kidneys, pancreas, and spleen unsuitable for transplantation were obtained via the UCSF Viable Tissue Acquisition Lab (VITAL) Core (IRB 20-31618). Kidneys were maintained on a Lifeport Kidney Transporter (Organ Recovery Systems) with circulating Viaspan solution for up to 36 hours after procurement. Pancreas and spleen were stored in University of Wisconsin (UW) solution at 4°C for up to 12 hours after procurement. All organs were delivered to our laboratory in static cold storage. Expired human allogenic packed red blood cells (RBCs) and fresh frozen plasma (FFP) were provided as a gift from the UCSF blood bank.

### Organ preparation and cannulation

In both porcine and human kidneys, excess perirenal tissue was removed using suture ligatures or Ligasure bipolar cautery (Covidien). The renal artery was identified and cannulated using an appropriately sized Luer lock tubing connector (Fig. S1). The cannula was secured using a single 2-0 silk tie (Ethicon). The arterial cannula was flushed with UW solution to help identify the renal vein. The vein was then similarly cannulated and secured with a silk tie. The artery was flushed a final time to ensure no additional vessels were missed. In the case of multiple arteries or veins, all vessels were cannulated and joined into a single inflow (arterial) and outflow (venous) line using Y connectors. The ureter was cannulated using a 5 Fr ureteral catheter and secured using a silk tie.

Human spleens were procured with the pancreatic tail intact to preserve long, single splenic vessels, including both the artery and vein. Excess peri-splenic and peri-pancreatic fatty tissue was ligated using silk ties. The splenic artery was identified and cannulated with a Luer lock tubing connector, secured using a 3-0 silk tie. The splenic vein, running parallel to the artery, was similarly cannulated and tied. The artery was flushed with UW solution, with efflux confirmed through the cannulated splenic vein. The inferior pole of the spleen, transected at procurement for standard donor HLA typing, was inspected for bleeding. Small vessels at the cut surface were ligated using silk ties and an Autosuture Premium Surgiclip II Clip Applier (Medtronic). In cases of significant bleeding during test perfusion with heparinized saline, the cut edge was imbricated with interrupted horizontal mattress sutures using 2-0 silk on a CT-1 needle (Ethicon). Human pancreata were procured using standard surgical techniques for transplantation, with en bloc resection of the pancreas, C-loop of duodenum, spleen, and associated small-bowel mesentery. Excess adipose tissue was excised, and the mesentery was divided between silk ties. The spleen was mobilized and detached between sequential silk ligatures. The splenic artery and superior mesenteric artery were dissected free, cannulated with appropriately sized Luer lock connectors, and secured with 3-0 silk ties. The two arterial inflows were then joined using a Y-connector. The portal vein was isolated and cannulated in a similar fashion. The sphincter of Oddi was identified, probed to confirm patency into the duodenum, and cannulated with a 5 Fr stent. The stapled ends of the duodenum were oversewn with a running 3-0 Prolene suture on an MH needle (Ethicon). The arterial system was flushed via the Y-connector with UW solution, and efflux was confirmed through the cannulated portal vein. Test perfusion with heparinized saline was then performed, and bleeding points were controlled using silk ties and an Autosuture Premium Surgiclip II Clip Applier (Medtronic).

### Perfusate composition

An autologous leuko-reduced blood preparation was used for porcine perfusion experiments as described above. For human experiments, packed RBCs were mixed with thawed FFP in a 1:1 ratio. All RBCs, FFP, and human donor organ were selected for serocompatibility.

Both porcine and human blood mixtures were supplemented with a perfusate additive consisting of Dextran 40 (Thermo Fisher), AlbuMAX lipid-rich bovine serum albumin (Gibco), sodium bicarbonate (7.5%), and calcium gluconate (Fresenius Kabi) osmotically balanced with Plasma-Lyte (Baxter) and sterile water. Nutrient supplementation (RPMI 1640 (Gibco) + GlutaMAX (Gibco) + dextrose (Duravet) + HumulinR (Lilly)) was infused via a syringe pump for perfusion experiments exceeding 6 hours duration.

### Physiologic monitoring during normothermic perfusion

Arterial pressure was monitored using a digital blood pressure analyzer (Digi-Med). Temperature was monitored using a digital infrared thermometer (ThermoBio). Arterial flow rate was measured using a clamp-on ultrasonic flow sensor (Sonotec). Blood gases and electrolytes were assessed using an i-STAT 1 blood gas analyzer (Abbott) with CG8+ cartridges. Ultrasonic assessment (General Electric) in grayscale and color Doppler was performed using a linear probe.

### Porcine nephrectomy and renal autotransplantation protocol

This large animal protocol was developed with University of California San Francisco and University of Colorado IACUC approval (AN203191, 01437, respectively). This protocol was adapted from methodology described by Kaths et al.^20^ Female juvenile Yorkshire pigs (∼40-50 kg) were brought to the facility and allowed to acclimate for at least 72 hours. A unilateral nephrectomy was performed under anesthesia through a midline incision. 5000 u heparin and 10 mg Lasix were administered prior to ligating the renal vessels. Upon removal, the kidney was quickly flushed with cold storage solution through the artery until the venous effluent turned clear.

In the immediate autotransplantation condition, autotransplantation of the removed kidney was performed without *ex vivo* perfusion. The kidney was prepared for autotransplantation on a back table. The infrarenal aorta and inferior vena cava (IVC) were exposed. Once the vessels were adequately mobilized and a suitable location for vascular anastomosis was identified, 5000 u heparin was given, and the IVC was occluded with a Satinsky vascular clamp. The IVC was incised and flushed with heparinized saline. The venous anastomosis was performed in a running fashion with 6-0 prolene suture. Upon completion of the anastomosis, a Debakey clamp was used to occlude the renal vein, and the IVC clamp was released. The aorta was then occluded with a Satinsky vascular clamp, and an aortotomy was made using a knife and 4.0 mm aortic punch (Medtronic). The arterial anastomosis was then performed also with 6-0 prolene suture. Upon completion, a second dose of 5000 u heparin was given and the vascular clamps were released. The ureter was then spatulated and anastomosed to the bladder with a running 5-0 polydioxanone (PDS) suture. Contralateral nephrectomy was then performed, and the bladder was emptied with a syringe. Abdominal closure was performed in three layers.

For autotransplantation experiments involving NEVKP, the animal was closed and allowed to recover for two days prior to autotransplantation. The removed kidney was perfused on our NEVKP device with leukoreduced autologous blood (in these cases, 500 mL autologous blood was collected during the nephrectomy) for 12 or 24 hours. Upon conclusion of NEVKP, the kidney was flushed with cold storage solution and placed in cold storage until autotransplantation (procedure as described above) on Day 0.

Following autotransplantation, the animal was recovered for two to 30 days. During this period, urine production was qualitatively assessed by placing absorbent pads beneath the cage away from the animal’s water source; these pads were checked twice daily for visible urine spots. Ultrasonic assessment (General Electric) of the autograft in grayscale and color Doppler was performed. Bladder urine and the autotransplanted kidney were harvested during necropsy and preserved for downstream analysis.

### Serum and urine chemistries

Serum and perfusate chemistries were performed on an iStat device (Abbott Medical) with GC8 and Chem8 cartridges. Urine chemistries were performed by Antech Diagnostics. Urinalysis was performed on a urine analyzer (McKesson).

### Measurement of glucose, c-peptide, and insulin

Arterial perfusate samples were analyzed using the i-STAT handheld blood analyzer (Abbott Point of Care, Princeton, NJ, USA) with CG8+ cartridges. Point-of-care measurements of glucose, pH, pO₂, pCO₂, sodium, and potassium were obtained at regular intervals to monitor oxygenation status and guide pH and electrolyte adjustments. Analyses were performed according to the manufacturer’s instructions, with cartridges maintained at room temperature prior to use. Baseline glucose values were recorded prior to glucose challenge. Glucose boluses were administered as D-glucose dissolved in lactated Ringer’s solution, and perfusate glucose was reassessed one hour post-challenge using the i-STAT analyzer. Concurrently, 5 mL perfusate samples were collected, centrifuged at 2000 rpm for 10 min, and the serum was separated, aliquoted, and stored at −20 °C for subsequent analysis. C-peptide and insulin concentrations were measured using Quantikine Human C-peptide and Insulin ELISA kits (R&D Systems) according to the manufacturer’s instructions. ELISA plates were read on a microplate reader, and data were analyzed using Microsoft Excel (Microsoft, Redmond, WA, USA) and GraphPad Prism (GraphPad Software, San Diego, CA, USA).

### Histology

Tissue samples were fixed in 10% formalin for 24-48 hours, stored in 70% ethanol, and then embedded in paraffin. Tissue sections sliced to 4 µm and mounted on positively charged Superfrost microscope slides (Fisher Scientific). Hematoxylin and eosin (H&E) staining was performed using a standard method.

### Flow cytometry

Arterial perfusate samples were collected at defined intervals in BD Vacutainer blood collection tubes containing K₂-EDTA (Fisher Scientific) and stored at 4 °C until processing. At the terminal time point, whole spleen was harvested, placed in saline, and maintained at 4 °C. Perfusate samples underwent red blood cell (RBC) lysis followed by Ficoll density gradient centrifugation to isolate lymphocytes, which were subsequently stained for flow cytometry using the Cytek cFluor Immunoprofiling 14-color RUO kit in combination with the ViaDye Red Fixable Viability Dye kit (Cytek Biosciences). Spleen tissue was digested to generate a single-cell suspension, subjected to RBC lysis, and stained using the same antibody panel. Flow cytometry was performed on a Cytek Aurora spectral flow cytometer (Cytek Biosciences), and data were acquired according to the manufacturer’s specifications.

### Statistical analysis

All quantitative data were analyzed using R Studio. Data are shown as mean ± standard deviation (if normally distributed) or mean and interquartile range (IQR) (25^th^ – 75^th^ percentile; if not normally distributed). Group comparisons were performed using Student’s t-test or Wilcoxon rank-sum. Statistical significance was set at p < 0.05.

## RESULTS

### An ultra-low cost NEVKP circuit can be constructed using commercially available components

We first constructed a simple organ perfusion circuit (Prototype A) using a peristaltic pump, a neonatal hollow fiber oxygenator, and Tygon tubing (Fig. 1A). All circuit components were sourced from commercial vendors for a total cost of less than 1500 US dollars (Table 1). A pressure sensor was connected to the arterial tubing to monitor the inflow pressure (Prototype A, segment I), and an ultrasonic flow sensor was attached to either the arterial (Prototype A, segment I) or venous tubing (Prototype A, segment B). Alternatively, venous flow rate was also measured by intermittently diverting the outflow into a graduated cylinder.

The three prototypes shown in Figure 1 represent the chronological evolution of our circuit design. In Prototype A, the circuit was designed according to previously described normothermic organ perfusion devices with the organ and oxygenator in series.^21^ A perfusate reservoir was fashioned from a clean plastic 1 L bottle, and an open plastic container was used to house the kidney. Normothermic (37 °C) warming was provided by a hot plate beneath the kidney and a water bath around the reservoir. This prototype allowed us to perform initial short-term kidney perfusion experiments.

In Prototype B, a bypass segment was added to help regulate perfusion pressure to accommodate kidneys that could not tolerate the minimum 100 mL/min of flow required by the oxygenator. The bypass segment thus served as an adjustable pressure release valve and the parallel design of the circuit allowed the overall flow through the oxygenator to be maintained. In Prototype B, pressure regulation via the bypass segment was accomplished by gravity. The flow was directed to a height of ∼100 cm above the kidney to achieve a pressure limit of ∼74 mmHg. Importantly, the fluid bag at the top of the bypass segment was left open to atmosphere to create this pressure differential.

Finally, Prototype C was designed to facilitate longer periods of perfusion (∼24 hrs). An infusion drip was added to the system to replenish nutrients and heparin; this could also be performed manually. To replace the open-to-air mechanism used in Prototype B, we incorporated a pinch valve using a segment of silicone tubing that could be compressed to varying degrees, essentially serving as an adjustable flow resistor. Perfusion pressures increased as the angle tightened (Fig. 1A, Prototype C, inset).

A sterile containment bag for the kidney was also added. This was fashioned from a 3 L fluid bag (Baxter), utilizing the drainage port at the bottom to allow for collection and recycling of venous hemorrhage. The kidney was stabilized within the bag using a series of neodymium magnets. Additional detail on bag design is provided below.

A reservoir was customized by drilling holes to accommodate drainage ports in a 300 mL neonatal suction canister or a T225 tissue culture flask. These were favored over the prefabricated Capiox FX05 reservoir due to simplicity of design and ease of cleaning for reuse. Multiple openings were drilled in the lid to allow all fluids to drain into the reservoir, including a port for urine recycling. Rather than warm the reservoir directly, perfusate warming was achieved by cycling ∼37 °C water through the warming ports of the oxygenator.

### Cost reduction during circuit development: practicing with non-heparinized porcine kidneys and reusing circuit components

To mitigate experimental costs during the early phases of the learning curve, disposable circuit components were cleaned and reused multiple times. After each experiment, all components were air dried after being flushed with Tergazyme, 10% bleach, and deionized water. In particular, we found that drying out the oxygenator between uses was the key to preserving oxygenator efficiency (Supplementary Fig. 1). This was accomplished by blowing dry air through both the gas and perfusate channels overnight. Using this approach, oxygenators could be used 3-5 times before they showed discoloration or wear.

To reduce the cost of organs and blood, fresh porcine kidneys and autologous blood were obtained during necropsy of other porcine experiments in our animal facility or as unutilized byproducts from a local slaughterhouse. In both cases, kidneys and blood were obtained free of charge. Autologous blood was heparinized immediately and stored in CPDA (citrate-phosphate-dextrose-adenine) buffer. To optimize organ viability, cold storage of kidneys was typically achieved within 20 minutes of circulatory arrest. These materials were then transported to our laboratory and prepared for NEVKP (Fig. 1B, see methods).

### Non-heparinized kidneys can be perfused *ex vivo* by maintaining a constant perfusion pressure

High intrarenal resistance was a frequently encountered challenge when attempting to perfuse porcine kidneys that were harvested without systemic heparinization prior to cardiac death. Despite liberally flushing each kidney with heparinized solution upon harvest, diffuse mottling of the renal parenchyma was observed, consistent the presence of microvascular clots (Fig. 2A). When perfusing these kidneys at the oxygenator’s minimum flow rate (100 mL/min), the arterial pressure would rise above 300 mmHg and cause hemorrhage around the hilum and into the collecting system (Fig. 2B). Histologic examination of these tissues after 1 hour of NEVKP under high pressures revealed distorted glomerular and tubular architecture and diffuse extravascular extravasation of red blood cells (Fig. 2D).

**Figure 2.**
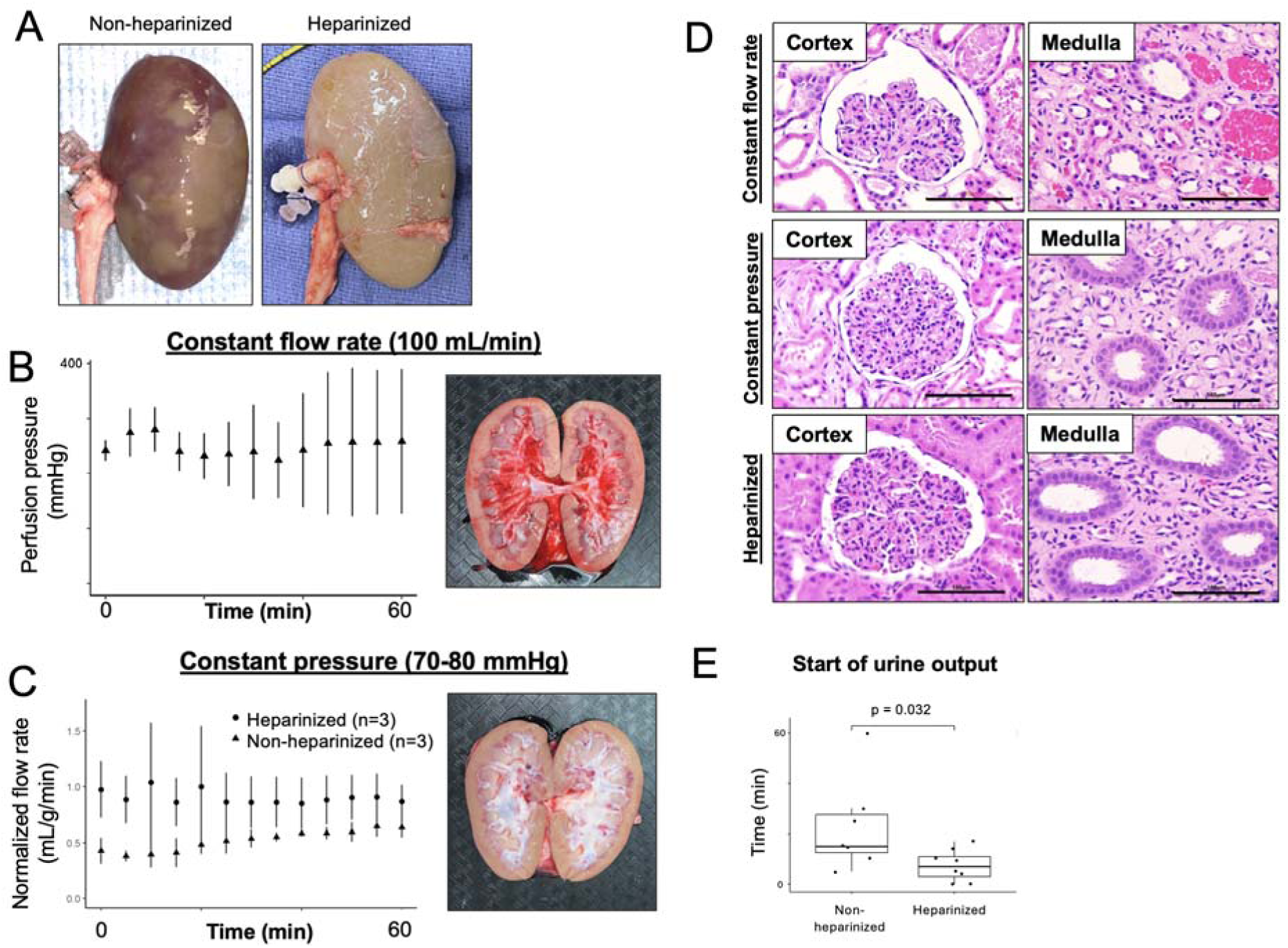
Perfusion of non-heparinized kidneys. A. Mottled appearance of a flushed kidney harvested from an animal without systemic heparinization compared to a flushed kidney from a heparinized animal. B. Perfusion pressure during first 60 minutes of NEVKP when flow rate was maintained at a constant 100 mL/min. Representative photo of pressure-related hemorrhage in a non-heparinized kidney. C. Normalized perfusion flow rate during first 60 minutes of NEVKP in heparinized and non-heparinized kidneys when perfusion pressure was held constant between 70 and 80 mmHg. Representative photo of non-heparinized kidney perfused for 60 minutes without hemorrhage. D. Representative H&E stains of porcine renal tissue according to perfusion condition. Scale bars = 100 µm. E. Comparison of urine production start times during NEVKP for heparinized and non-heparinized kidneys.

**Figure 3.**
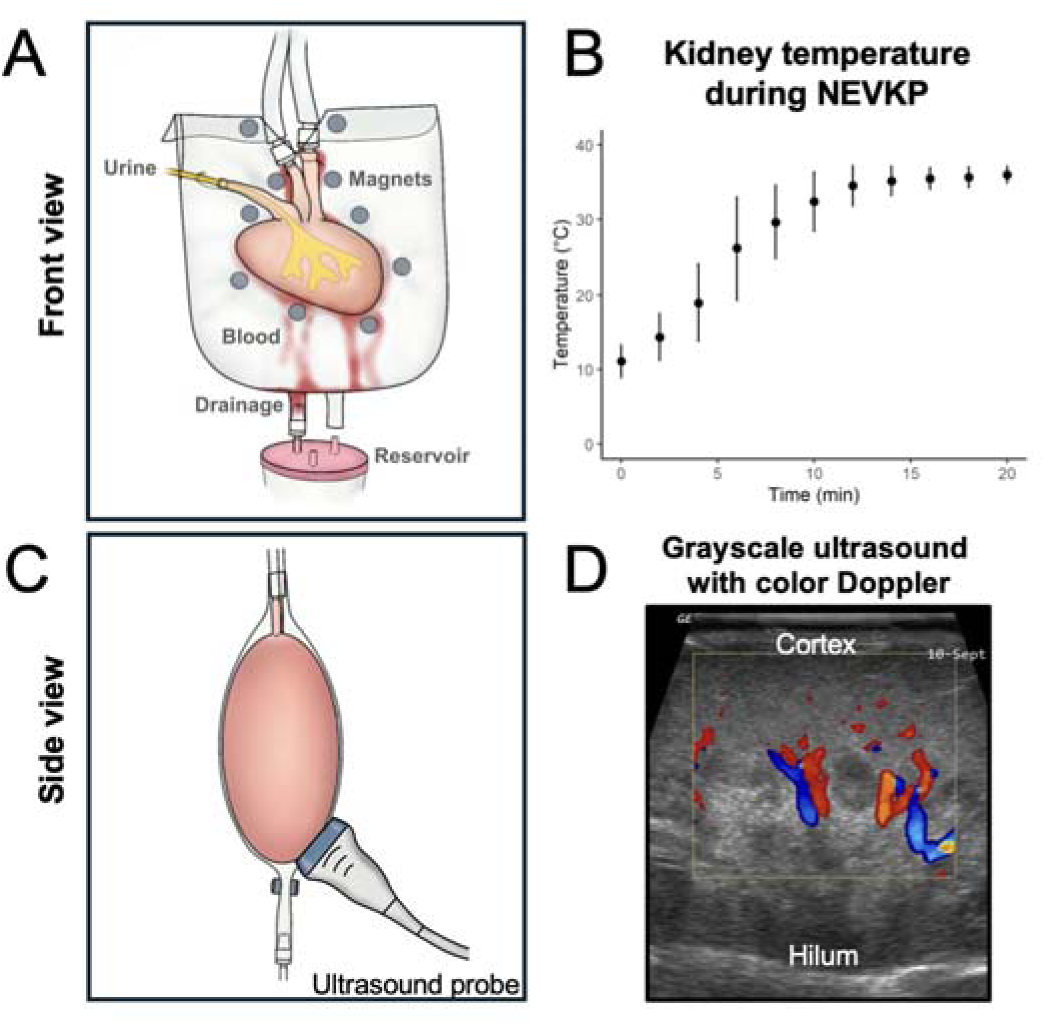
Normothermic *ex vivo* kidney containment bag. A. Front view of bag design. B. Surface temperature of kidney during NEVKP during first 20 minutes (n=4). C. Side cross-sectional view of kidney within the bag with ultrasound probe. D. Representative grayscale ultrasound with color Doppler image of kidney during NEVKP.

To overcome this problem, we incorporated the pressure-regulating bypass mechanisms described above (Prototype B and C) to permit lower flow rates to the kidney (< 100 mL/min) while maintaining a higher flow rate through the oxygenator (≥ 100 mL/min). While holding the perfusion pressure constant, we observed a gradual increase in flow to non-heparinized kidney (Fig. 4C) and a resolution in the mottled appearance of the kidney, likely consistent with the dissolution of microvascular clots. After 60 minutes of perfusion, these kidneys were histologically indistinguishable from heparinized kidneys (Fig. 2D). From a functional perspective, the primary consequence of using non-heparinized kidneys was a delayed onset of urine production (22.9 (10-30) mins vs 7.4 (3-11) mins, p = 0.03) (Fig. 2E). While systemic heparinization prior to organ harvest remains ideal, this approach allowed us to progress in our learning curve using non-heparinized organs (Supplementary Fig. 2).

**Figure 4.**
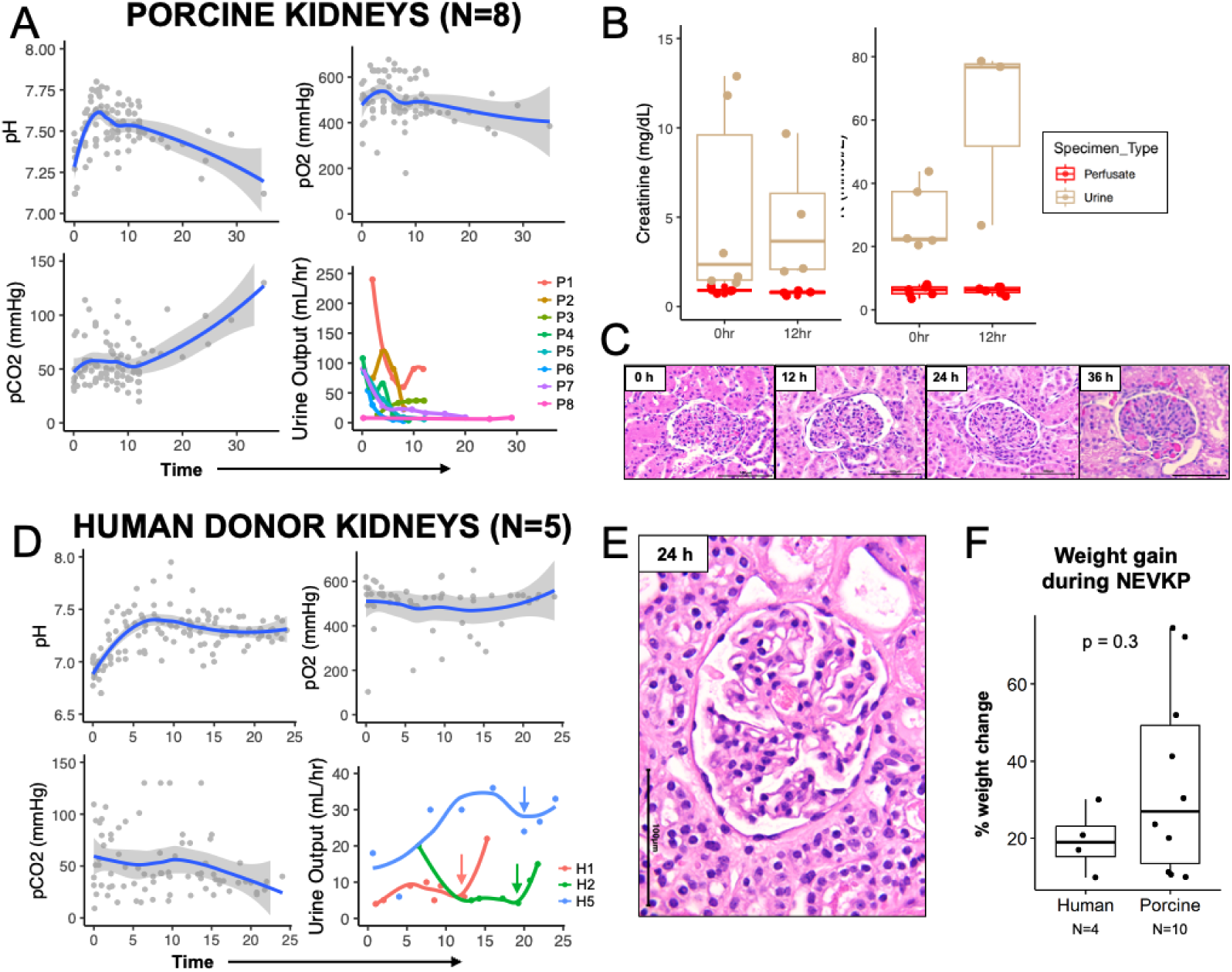
Prolonged NEVKP of porcine and human kidneys. A. Aggregate pH, pO2, pCO2, and urine output over time from eight porcine NEVKP experiments. B. Perfusate and urine creatinine and potassium concentration at 0 and 12 hrs. C. Representative H&E stain of glomerulus and renal tubules between 0-36 hours of NEVKP. D. Aggregate pH, pO2, pCO2, and urine output over time from five human donor NEVKP experiments. E. Representative H&E stain of glomerulus and renal tubules of human donor kidney after 24 hours of NEVKP. F. NEVKP-associated weight gain of human and porcine kidneys.

### A containment bag with adjustable magnets provides stability and perfusate recycling during prolonged NEVKP

Most NEVOP systems house the organ in a rigid container. Using this approach in prototypes A and B, we found that frequent adjustment of the vascular cannulas was required to prevent kinking and twisting. The venous tubing was particularly prone to occlusion due to the flimsy nature of the porcine renal vein. The need for constant manipulation was labor-intensive and raised concerns for maintaining sterility during prolonged NEVKP experiments. Separately, we also recognized the need to recycle persistent leakage of perfusate. Venous hemorrhage around the hilum was unavoidable despite our best efforts to ligate small vessels using suture ligatures or bipolar cautery.

To address these needs, the kidney was housed in an empty 3L bag (Fig. 3A). Within the bag, the kidney and cannulas were fixed in space using a series of neodymium magnets. The tubing exited at the top and was further secured to a stand. This configuration minimized the frequency of kinking or twisting at the hilum, and adjustments could be performed without opening the bag.

To enable perfusate recycling, the magnets were spaced 1-2 cm apart to allow leaked perfusate to drain around the kidney without pooling. The drainage port at the bottom of the bag was connected to the reservoir, allowing the perfusate to be promptly recycled. We found that the native drainage port could accommodate a maximum flow rate of 600.5 ± 42 mL/min, which was useful in cases where the renal vein was not cannulated and instead allowed to bleed freely into the bag.

As the bag was not directly warmed by a heating element, we assessed whether normothermia could be maintained by warming the perfusate alone by running heated water through the oxygenator. We observed that the temperature of kidneys reached 37 °C within 20 minutes of perfusion initiation and could be maintained throughout the duration of NEVKP (Fig. 3B).

Finally, we noted that having an airtight contact between the bag and the kidney provided the additional benefit of facilitating ultrasonography during NEVKP without impacting sterility (Fig. 3C). Perfusion of different regions could be assessed in real-time using color Doppler (Fig. 3D).

### NEVKP of porcine and human kidneys can be achieved for up to 24 hours using prototype C

Having overcome the initial challenges of establishing an NEVKP system, we next sought to explore whether prolonged NEVKP (≥ 24 hours) could be achieved using prototype C. For these experiments, the peristaltic pump was replaced with a centrifugal pump (Terumo-Sarns Delphin II) to reduce hemolysis during longer perfusion times.^19^

Eight porcine kidneys were perfused: six for 12 hours (P1-P6), one for 24 hours (P7), and one for 36 hours (P8). In each experiment there was a pattern of initial alkalosis peaking at pH 7.75 between hour 4-5 of NEVKP (Fig. 4A); this was typically managed by reducing the flow rate of carbogen to the oxygenator or by slowly adding up to 10 mL of 0.1M HCl to the perfusate in drop-wise fashion. Throughout all experiments, pO2 remained above 400 mmHg, indicating adequate oxygenation, while pCO2 gradually increased over time during the 24 and 36 hour experiments.

Urine production varied widely within and across experiments (Fig. 4A). Creatinine and potassium concentrations were much higher in the urine compared to the perfusate at 0 and 12 hrs, indicating active clearance. To further explore renal potassium handling during NEVKP, urine recycling was not performed for P8, which was allowed to run for 36 hours. In this experiment, the concentration of potassium was initially high (likely due to hemolysis during blood collection and processing) but reached homeostasis between 3.2 and 4.7 mmol/L at hour 5 of NEVKP through the remainder of NEVKP (Supplementary Fig. 3). This trend corresponded to a peak in urinary potassium concentration at hour 5 followed by a gradual reduction over time.

Histologic examination over 36 hours of NEVKP revealed grossly intact architecture and cellular structure (Fig. 4C). There was evidence of congestion and hyaline deposits in the glomerulus at 36 hours along with evidence of osmotic tubular changes starting at 24 hours.

Prolonged NEVKP was also performed in four non-transplantable human donor kidneys from four donors. Donor demographics and clinical profiles are shown in supplementary table 1. These kidneys were perfused for 18 hrs (H1), 24 hrs (H2), 16 hrs (H3-4), and 24 hrs (H5). H1 was terminated for inadequate oxygenation due excess condensation in the oxygenator; pO2 and pCO2 were otherwise maintained throughout perfusion for the others (Fig. 4D). Physiologically, the pH stabilized around 7.3-7.4 at around the five-hour mark in each experiment (Fig. 4D). Urine output H1, H2, and H5 slowly decreased throughout NEVKP but increased with administration of furosemide (Fig. 4D). H3 and H4 exhibited brisk gross hematuria from traumatic catheterization; as a result, urine output could not be accurately measured.

Histologically, the perfused human kidneys appeared similar to porcine kidneys at 24 hours with evidence of osmotic tubular changes (Fig. 4E). Both human and porcine kidneys exhibited weight gain during NEVKP, consistent with gross edema (Fig. 4F).

Interestingly, H1 had an unperfused segment due to an accessory artery that we ligated prior to NEVKP, providing an internal unperfused control for comparison (Supplementary Fig. 4A). The tissue architecture of the perfused segment was preserved compared to the unperfused segment, which exhibited signs of necrosis (Supplementary Fig. 4B).

Taken together, these data indicate that while further perfusate optimization is needed, human and porcine kidneys exhibit evidence of viability and function during prolonged NEVKP with prototype C.

### Porcine kidneys function *in vivo* after prolonged NEVKP

To test the *in vivo* performance of kidneys after NEVKP, we adapted a porcine renal autotransplantation protocol using juvenile Yorkshire pigs (40-50kg) based on previously described methodology.^22^ We compared autotransplantation outcomes among three NEVKP conditions: no NEVKP (i.e. nephrectomy with immediate autotransplantation), after 12 hours of NEVKP, and after 24 hours of NEVKP. In each condition, the contralateral kidney was removed concurrently at the time of autotransplantation to render the animal dependent on the autograft for urine production (Fig. 5A).

**Figure 5.**
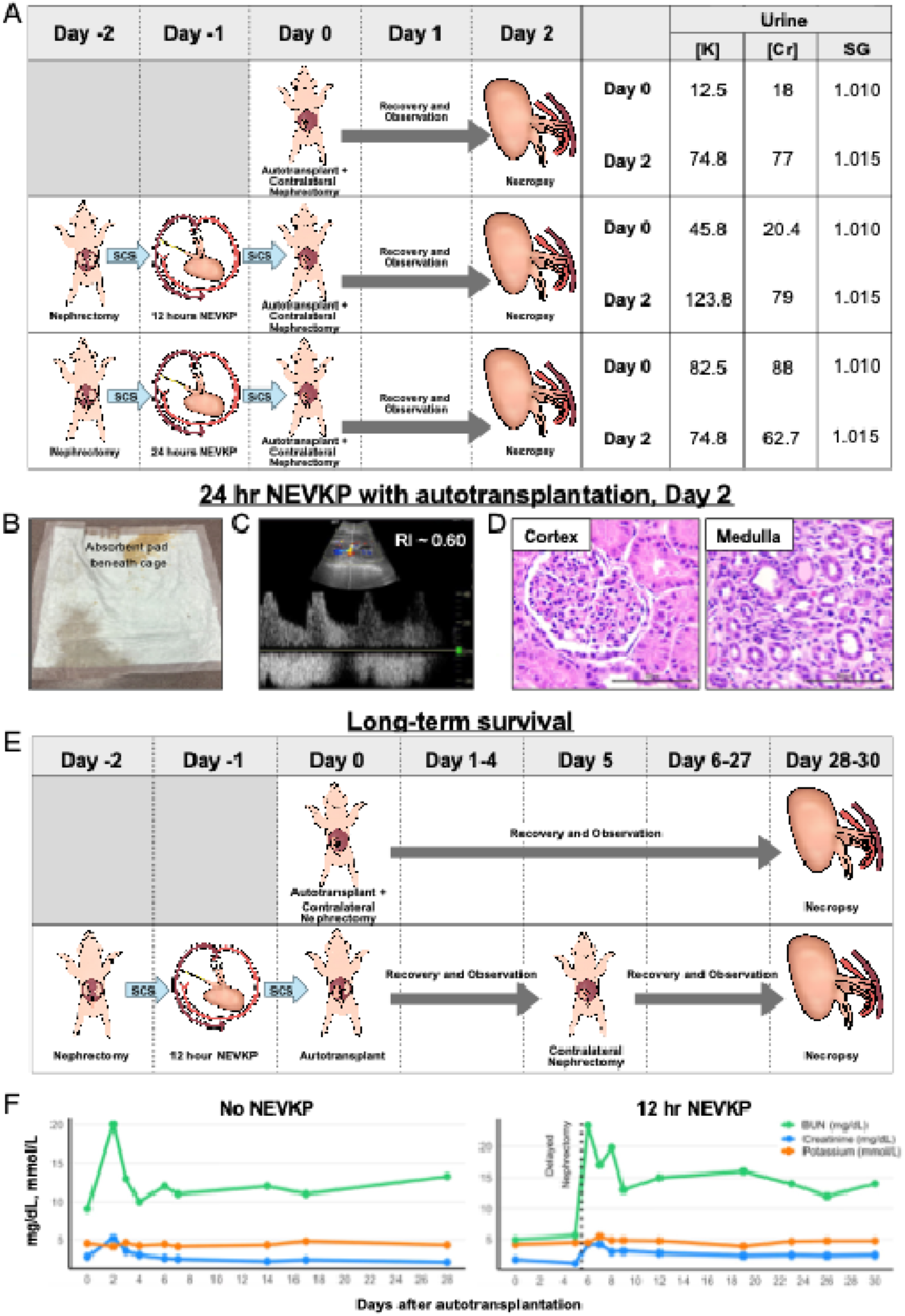
Autotransplantation of porcine kidneys after NEVKP. A. Experimental schematic of NEVKP and autotransplantation experiments with comparison of urinary potassium (mmol/L), creatinine (mg/dL), and specific gravity. B. Representative photo of an absorbent pad placed beneath the animal’s cage with a wet area (lower left) indicative of urine, taken on Day 2 after autotransplantation of a 24-hour NEVKP kidney. C. Representative image of Doppler waveform and resistive index of the 24-hour NEVKP autograft on Day 2 after autotransplantation. D. Representative H&E stain of 24-hour autograft taken on necropsy. E. Schematic of long-term survival experiment. F. Serum blood urea nitrogen (BUN), creatinine (Cr), and potassium (K) after autotransplantation. Scale bars = 100 µm. SCS = Static Cold Storage, RI = Resistive Index.

*In vivo* urine production was observed in each case. Urine production was noted upon completion of the vascular anastomosis in the case of the immediately autotransplanted kidney and the 12-hour NEVKP kidney. These animals urinated overnight between Day 0 and Day 1. Although immediate urine production was not observed for the 24-hour NEVKP kidney, the animal began to urinate between Days 1 and 2 (Fig. 5B). Doppler assessment of the autograft demonstrated adequate flow with a resistive index of ∼0.60 (Fib. 5C). All animals were noted to have at least 100 mL of urine in the bladder upon necropsy. In each case, the bladder urine contained elevated potassium and creatinine concentrations and had a specific gravity consistent with ongoing electrolyte clearance and urine concentration (Fig. 5A). Histologic examination of the kidney perfused for 24 hours demonstrated preserved architecture in both cortex and medulla and no evidence of osmotic tubular changes (Fig. 5D).

To assess whether a perfused kidney could sustain life, we assessed 4-week survival outcomes after renal autotransplantation after 12-hour NEVKP compared to autotransplantation without NEVKP. In the NEVKP condition, contralateral nephrectomy was performed 5 days after autotransplantation to mitigate the short-term risk of delayed graft function (Fig. 5E). Both animals survived for 4 weeks after autotransplantation with stabilization of serum blood urea nitrogen (BUN), creatinine (Cr), and potassium (Fig. 5F).

While more studies are needed to optimize the results, these experiments confirm that kidney viability and urine production are maintained during prolonged NEVKP.

### NEVKP system can be adapted to human pancreas and spleen

Having demonstrated that our NEVKP system supports renal viability and function, the platform was adapted to other solid organs, specifically the human pancreas and spleen. Non-transplantable human pancreata and spleens were obtained through the UCSF VITAL Core biobank. Minimal modifications to the containment bag were required to securely accommodate either organ while maintaining stable perfusion and secure cannulation.

The human donor pancreas was received attached to a portion of duodenum and small bowel mesentery. Dual arterial inflow was established via the splenic and superior mesenteric arteries, joined using a Y-connector, with venous outflow through the portal vein (Fig. 6A-B). Perfusion was conducted under normothermic conditions with mean arterial pressures (MAP) of 60–80 mmHg for 8 hours. To assess endocrine pancreatic function, glucose challenges were performed, and we observed progressive increases in C-peptide and insulin concentrations corresponding to rising glucose levels (Fig. 6C-E). These findings demonstrated that the NEVOP system supports both glucose sensing and hormone secretion in the ex vivo human pancreas.

**Figure 6.**
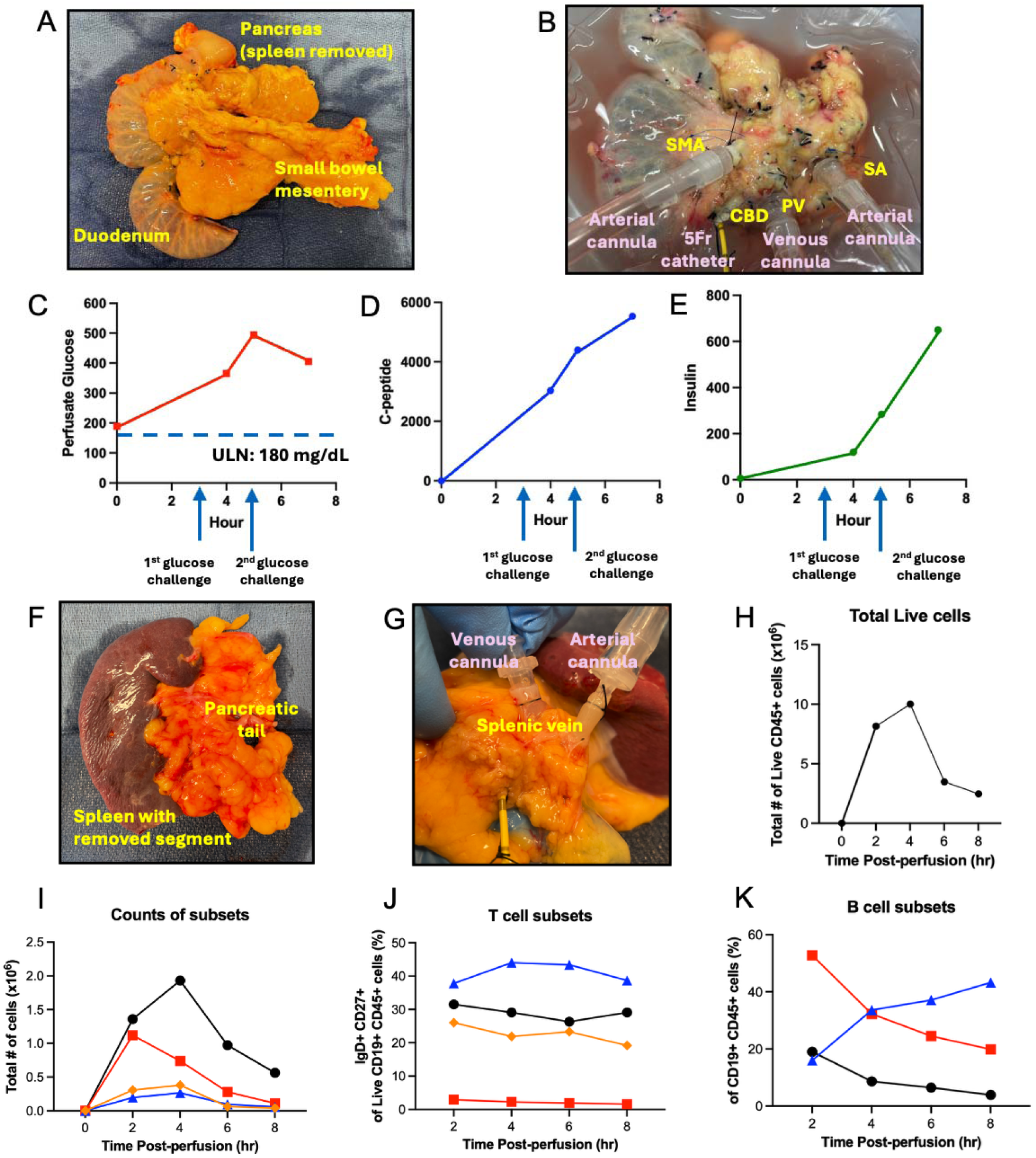
Normothermic ex vivo perfusion of human pancreas and spleen for functional and immunological assessment. A. Procurement of an adult human pancreas with attached duodenum and small bowel mesentery. B. Cannulation strategy for pancreas perfusion with arterial inflow via the superior mesenteric artery (SMA) and splenic artery (SA), venous outflow via the portal vein (PV), and drainage of pancreatic secretions through the common bile duct (CBD) / sphincter of Oddi. C-E. Functional assessment of perfusate glucose (C), C-peptide (D), and Insulin (E) concentrations indicating pancreatic endocrine activity upon glucose challenge. F. Procurement of an adult human spleen with attached pancreatic tail and intact splenic artery and vein. The inferior pole of the spleen was excised for HLA typing and immunologic testing. G. Cannulation of the spleen with inflow via the splenic artery, outflow via the splenic vein, and ductal stenting of the attached pancreatic tail. H. Flow cyometric analysis of live CD45+ leukocytes in the circuit perfusate over time. I. Kinetics of lymphocyte egress from the spleen during perfusion. J. Temporal dynamics of T cell subset release into the perfusate. K. Temporal dynamics of B cell subset release into the perfusate.

For spleen perfusion, arterial inflow was established via the splenic artery, with venous outflow through the splenic vein (Fig. 6F-G). Perfusion parameters mirrored those used for the pancreas (37 °C, MAP 60–80 mmHg, 8-hour duration). Splenic function and viability was gauged by the repopulation of immune cells in the leukoreduced perfusate over time. After confirming the absence of circulating lymphocytes at time 0, we detected an increase in the presence of live lymphocytes in the perfusate at subsequent time points, with preferential egress of specific T and B cell subpopulations into circulation. At the conclusion of perfusion, a wedge section of the spleen was processed into a single-cell suspension and analyzed by flow cytometry, confirming the persistence of viable lymphocytes within the tissue. These results indicate that the NEVOP system preserves both cellular viability and physiologic immune trafficking in the human spleen *ex vivo*.

Collectively, these studies demonstrate that the NEVKP system can be adapted with minimal modification to support *ex vivo* perfusion of human pancreas and spleen. The platform allows functional assessment of pancreatic endocrine activity and real-time monitoring of spleen immune cell dynamics under near-physiologic conditions, providing a versatile tool for investigating organ-specific function, viability, and physiology in a controlled ex-vivo setting.

## DISCUSSION

Here we describe the creation of an ultra-low cost, open-source NEVOP platform for the prolonged perfusion of porcine and human kidneys. A pragmatic engineering approach was employed and achieved proficiency using nonstandard sources of porcine kidneys and blood to overcome the learning curve. A porcine renal autotransplantation model confirmed that *in vivo* kidney viability and function can be maintained on our platform for up to 24 hours. Other small solid organs such as pancreas and spleen also can be perfused with minor modifications. Several innovations were introduced to facilitate the technical aspects of small organ perfusion; this reproducible strategy to perform organ perfusion experiments is sustainable, can be performed in a humane fashion and can be used at scale.

Our approach makes normothermic *ex vivo* organ perfusion more accessible to the broader scientific community. Many published NEVKP studies rely on commercial organ perfusion systems—often facilitated by preexisting industry partnerships—or on cardiopulmonary bypass systems, options that are not readily accessible. The cost of acquiring or leasing these systems without institutional or corporate affiliations can be prohibitive. Our perfusion circuit represents an economically favorable alternative as it can be assembled for approximately $1,000–$1,500 using widely available materials. The modular design also allows for customization and iterative improvements based on specific research needs, an advantage not easily achieved with proprietary systems. By lowering the financial and technical barriers to NEVKP, this approach has the potential to expand access to organ perfusion research and accelerate innovation in the field.

Importantly, our approach also lowers the technical barrier to performing NEVKP. Our circuit is simple to construct, and its essential components do not require an extensive technological background to operate. The design of the bag to stabilize the vascular cannulas and recycle venous hemorrhage obviates the need for advanced suturing skills or painstaking attention to hemostasis during kidney preparation. The only surgical skills necessary to prepare a kidney for NEVKP are basic tissue dissection to harvest the kidney and simple knot-tying to secure the cannulas. These skills can be acquired easily with practice; many experiments can be performed by non-medically trained personnel.

A key innovation of our system is the integration of a pressure regulation system to accommodate kidneys that require lower perfusion flow rates while simultaneously satisfying the minimum flow rate of the oxygenator. By facilitating the perfusion of nonheparinized kidneys for NEVKP studies, organs can be obtained from a variety of sources to lower the cost. In our case, we obtained kidneys and blood via necropsy within our animal facility and from a local slaughterhouse to overcome the early learning curve. Our circuit design can also facilitate the perfusion of other small and delicate organs. In addition to human pancreas and spleen, which we have shown in this manuscript, we anticipate that other body parts such small intestine segments, gonads, limbs, and skin flaps can be perfused in a similar fashion. Virtually any organ with arterial inflow could be perfused on this system as long as there is a means for stable cannulation. This can open the door to interesting biological studies examine organ biology in insolation.

Although we have demonstrated that our system can achieve prolonged NEVKP in both porcine and human organs, there remain several important limitations. The materials and components used in our circuit may not be optimized for hemo- or biocompatibility. Additionally, despite meticulous attention to maintaining a sterile, closed environment during prolonged perfusion experiments, contamination could pose a challenge for longer periods of NEVKP. These issues will be addressed in future refinments of the system. There were also a number of experimental variables that were difficult to control for such as blood product storage duration, warm ischemia time, and pre-existing kidney injury—these factors almost certainly influenced NEVKP performance and contributed to the variability in results. Finally, we acknowledge that there remain numerous gaps in understanding for how to optimally perform NEVKP. For example, there are currently no evidence-based strategies to manage nutrition, osmotic balance, blood product exchange timing, and waste clearance *ex vivo*. The reproducibility and sustainability of our NEVKP system should enable rigorous large-scale experiments to investigate these important questions.

Ultimately, organ perfusion is an emerging technology that promises to unlock new avenues of discovery. While its most salient application lies in organ preservation for transplantation, additional uses include the development of therapeutics using *ex vivo* models and the *ex vivo* administration of organ-targeted therapies. The ability to perform *ex vivo* organ perfusion sustainably and at scale represents a powerful new research paradigm and should be made more accessible to the scientific community. As organ perfusion research programs advance at our respective institutions, our experience can be used as a blueprint to attract other research teams and expand the scope of this work.

## Acknowledgements

We would like to acknowledge the following individuals and organizations for generously providing materials, expertise, and/or feedback critical to the success of this project: Bob Bartlett and Alvaro Rojas-Pena (University of Michigan Extracorporeal Life Support Lab); Grace Hwang (Children’s Hospital of Philadelphia); Dan Crompton (Vitara Biomedical); Rachel Jones (UCSF Perioperative Services); Sylvie Baudart and Anja Strehlow (UCSF Department of Mechanical Circulatory Support); Clinton Jones (UCSF Department of Pediatric Perfusion); Joseph Akin (UCSF Blood Bank); Rizwana Saleem (American Red Cross); Allison Harf (Medtronic). We would also like to thank the veterinary staff of the UCSF Laboratory Animal Research Center and the University of Colorado Office of Laboratory Animal Research. Finally, we would like to thank Ahmad Salehi of Donor Network West with assistance with VITAL Core samples, and we deeply thank the organ and tissue donors and their families for their incredible generosity in helping to advance scientific research.

## Author Contributions

Conceptualization - HY, AF, JG, MS

Data collection and analysis - HY, NH, SC, JL, NM

Technical and logistical execution – HY, NH, SC, JL, NM, KH, RF, TS, JM, MV, WS, HB, JD, JE

Drafting of initial manuscript - HY, NH, SC, JL, NM

Supervision - MS, HY, AF, SR, JG, TC

## SUPPLEMENTARY MATERIALS

**Supplementary Figure 1.**
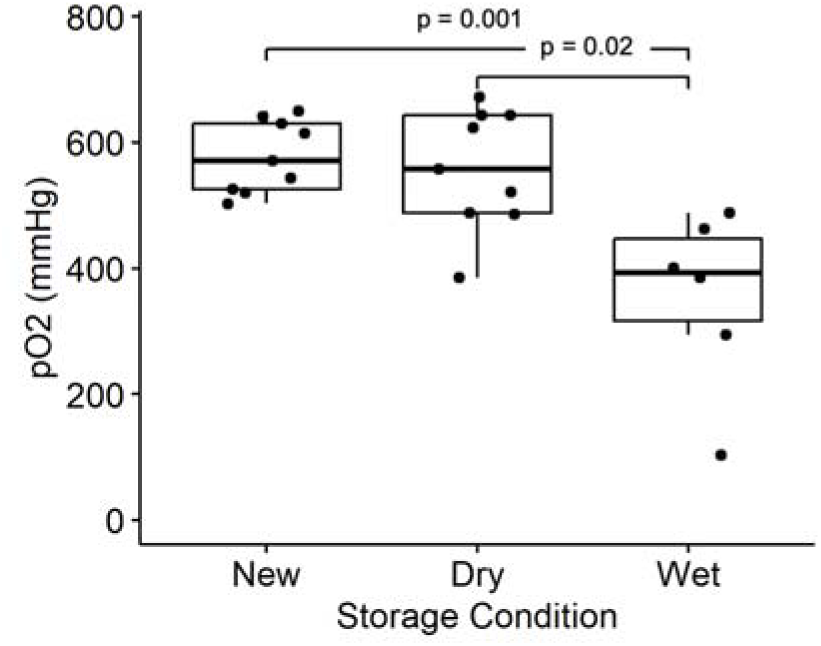
Perfusate pO_2_, an indicator of oxygenation efficiency, within first 15 minutes of NEVKP for new and used oxygenators.

**Supplementary Figure 2.**
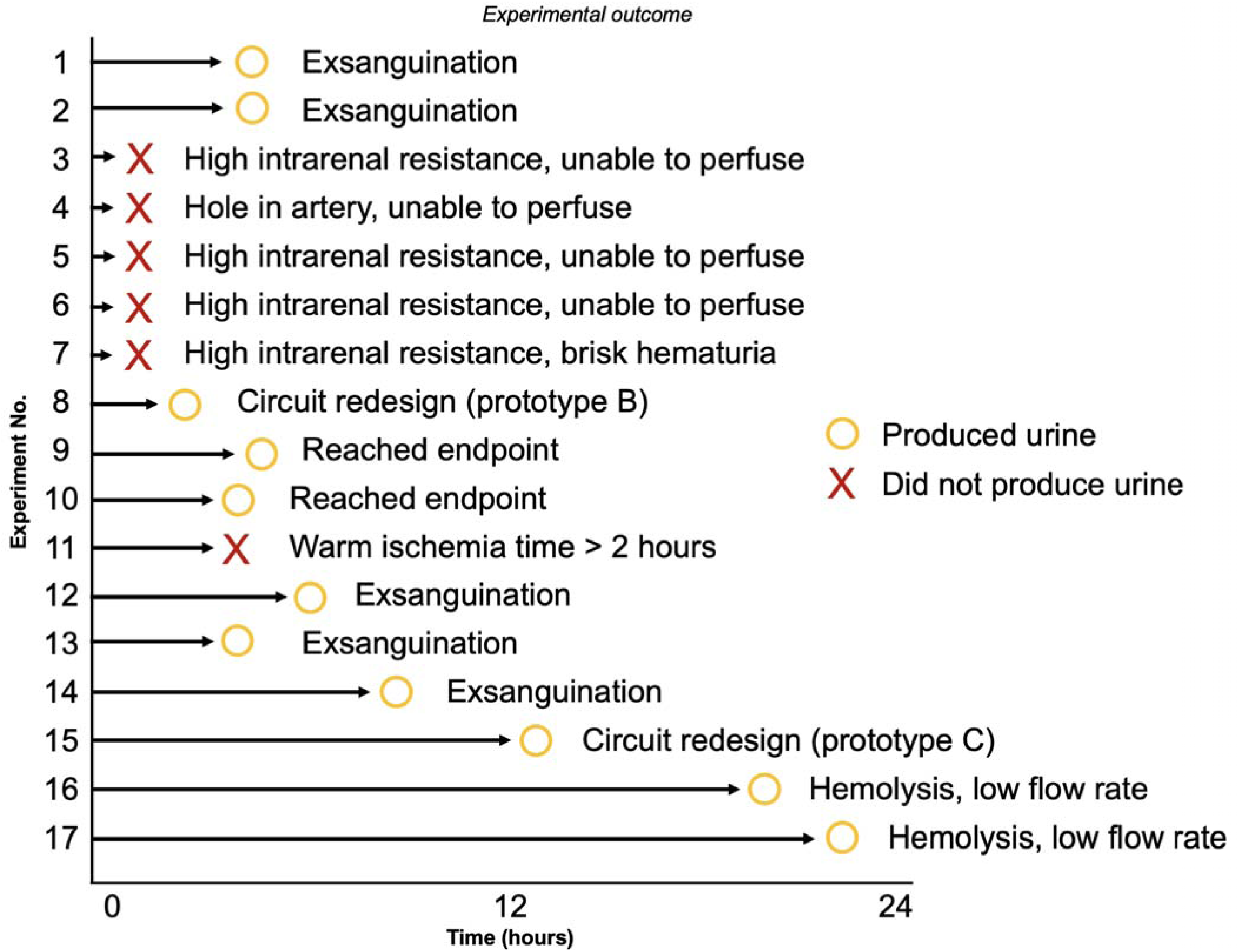
Learning curve for NEVKP. Experimental duration, outcomes, and comments for first 17 porcine NEVKP experiments.

**Supplementary Figure 3.**
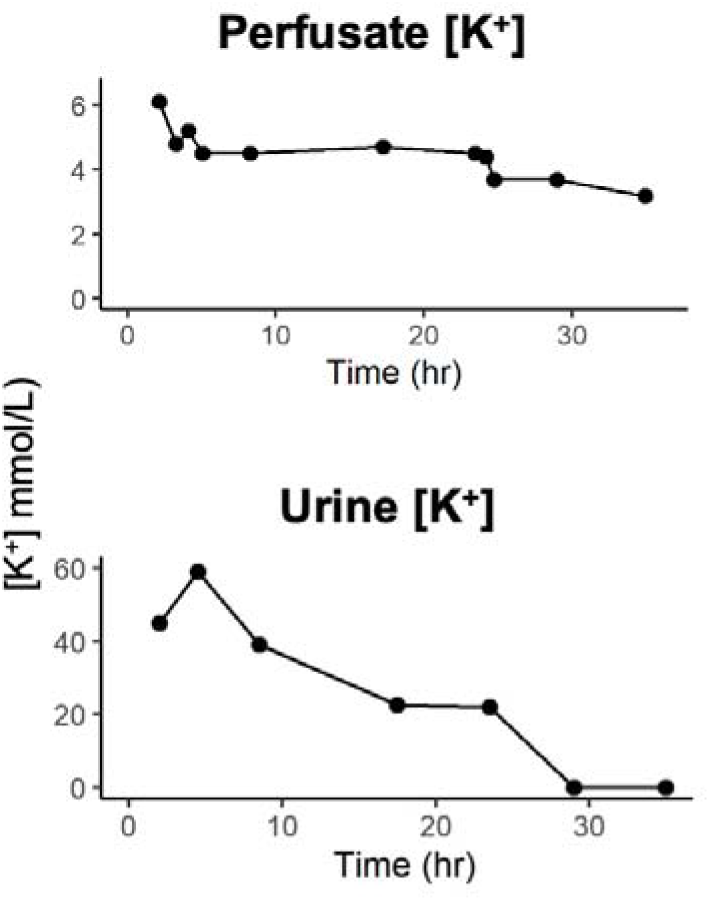
Perfusate and urine potassium concentration during 36-hour porcine kidney perfusion.

**Supplementary Figure 4.**
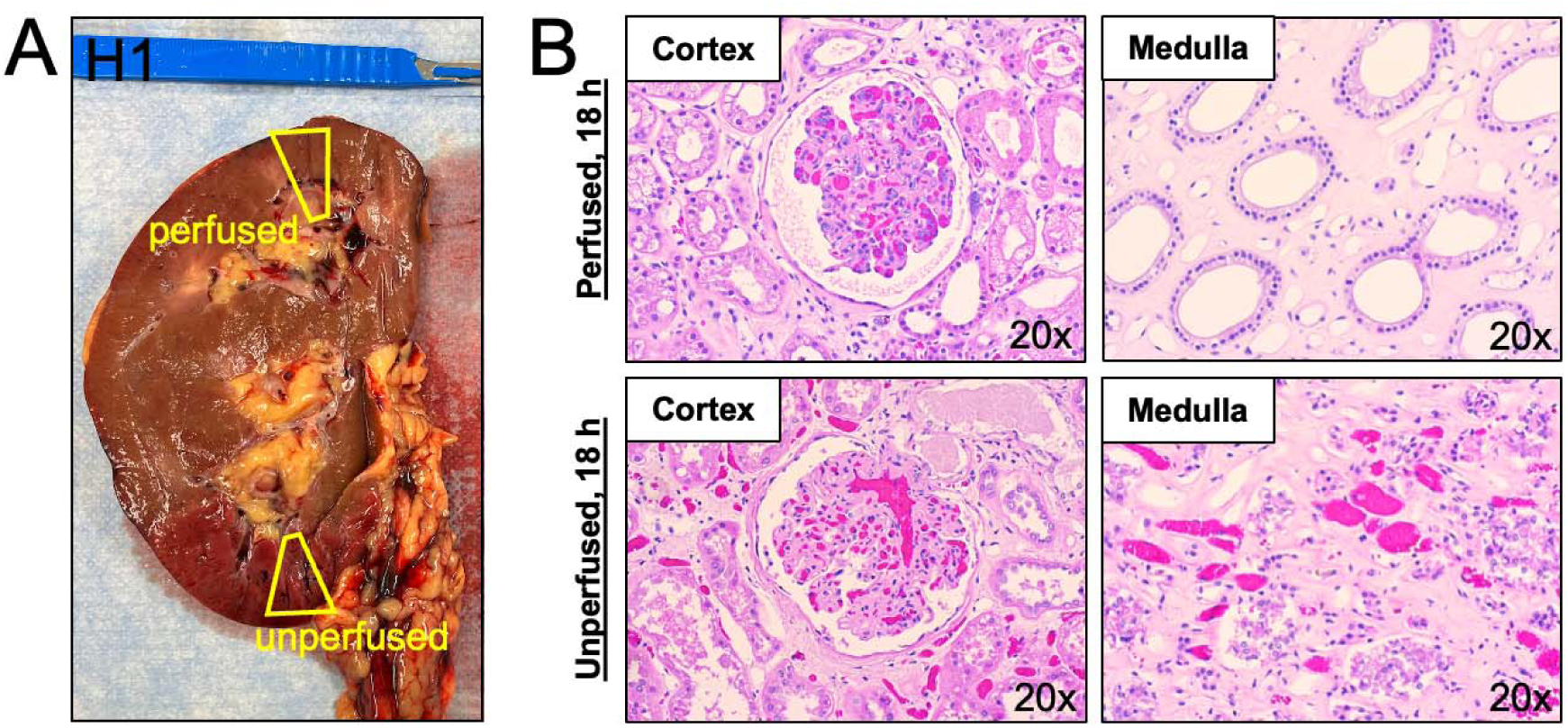
Human donor kidney perfused for 18 hours. A. Gross examination of kidney following perfusion. A small unperfused segment was noted in the lower pole due to ligation of an accessory artery prior to NEVKP. Yellow quadrangles indicate areas taken for histological examination. B. Representative histological images of cortex and medulla from perfused and unperfused areas.

**Supplementary Table 1.** Donor profile for untransplanted human kidneys. Cr = Creatinine. KDPI = Kidney Donor Profile Index. PMH = Past medical history.

| Kidney ID | Sex | Donor Age Range | Last serum Cr of donor | KDPI | Cold storage time | Significant PMH | Cause of death |
| --- | --- | --- | --- | --- | --- | --- | --- |
| H1 | M | 60-70 | 0.98 | 93% | 24-48 h | Hepatorenal syndrome | Respiratory |
| H2 | F | 60-70 | 0.89 | 90% | 24-48 h | Unknown | Anoxia (Seizure) |
| H3 (left),<br>H4 (right) | M | 60-70 | 2.22 | 85% | 24-48 h | Unknown | Anoxia (Cardiovascular) |
| H5 | F | 40-50 | N/A | 33% | 24-48 h | Heavy alcohol use | Anoxia (brain death) |

